# In-depth characterization of layer 5 output neurons of the primary somatosensory cortex innervating the mouse dorsal spinal cord

**DOI:** 10.1101/2020.04.02.021311

**Authors:** N. Frezel, E. Platonova, F.F. Voigt, J.M. Mateos, R. Kastli, U. Ziegler, T. Karayannis, F. Helmchen, H. Wildner, H.U. Zeilhofer

**Author notes:** corresponding authors: Dr. H.U. Zeilhofer & Dr. H. Wildner, Institute of Pharmacology and Toxicology, University of Zurich, Winterthurerstrasse 190, CH-8057 Zürich, Switzerland.

## Abstract

Neuronal circuits of the spinal dorsal horn integrate sensory information from the periphery with inhibitory and facilitating input from higher CNS areas. Most previous work focused on projections descending from the hindbrain. Less is known about inputs descending from the cerebral cortex. Here, we identified cholecystokinin (CCK) positive layer 5 pyramidal neurons of the primary somatosensory cortex (CCK^+^ S1-CST neurons) as a major source of input to the spinal dorsal horn. We combined intersectional genetics and virus-mediated gene transfer to characterize CCK^+^ S1-CST neurons and to define their presynaptic input and postsynaptic target neurons. We found that S1-CST neurons constitute a heterogeneous population that can be subdivided into distinct molecular subgroups. Rabies-based retrograde tracing revealed monosynaptic input from layer 2/3 pyramidal neurons, from parvalbumin (PV) positive cortical interneurons, and from thalamic relay neurons in the ventral posterolateral nucleus. WGA-based anterograde tracing identified postsynaptic target neurons in dorsal horn laminae III and IV. About 60% of these neurons were inhibitory and about 60% of all spinal target neurons expressed the transcription factor c-Maf. The heterogeneous nature of both S1-CST neurons and their spinal targets suggest complex roles in the fine-tuning of sensory processing.

## Introduction

In addition to the hindbrain, the cerebral cortex is a major source of descending input to the spinal cord (Lemon RN and J Griffiths 2005; Abraira VE et al. 2017; Wang X et al. 2017; Liu Y et al. 2018; Ueno M et al. 2018). Layer 5 pyramidal neurons of several cortical areas project to this site, including neurons residing in the motor and premotor cortices as well as in the somatosensory cortex (S1). In rodents as well as in humans, the axons of the corticospinal (CST) neurons travel through the internal capsule in the forebrain to enter the cerebral peduncles at the base of the midbrain. They then pass through the brainstem to form the pyramids, at the base of the medulla. The vast majority of the fibres decussate at this level to enter the spinal cord. From there, the axons of the rodent corticospinal tract (CST) run in the ventral part of the dorsal funiculus, while in humans the tract is located in the lateral white matter.

Most studies on the function of the CST have focused on fine motor control (Wang X et al. 2017; Ueno M et al. 2018) often in the context of spinal cord injury and spinal cord repair (Bareyre FM et al. 2005; Steward O et al. 2008; Fry EJ et al. 2010; Porrero C et al. 2010; Jin D et al. 2015). These studies have targeted either the whole CST, or CST neurons of the motor cortex. The presence of direct synaptic contacts between CST neurons that descend from S1 and spinal interneurons (Abraira VE et al. 2017; Liu Y et al. 2018; Ueno M et al. 2018) suggests that CST neurons also play an important role in somatosensory processing, beyond sensorimotor integration. This is in line with previous findings that CST neurons from the motor (M1) and S1 cortices contact distinct populations of spinal interneurons (Ueno M et al. 2018). The functional analysis of specific parts of the CST has been in part limited by the lack of tools to specifically target defined subgroups of CST neurons (e.g., CST neurons that descend from a defined cortex area to a specific spinal cord region). Transgenic mouse lines (Bareyre FM et al. 2005; Porrero C et al. 2010) and virus-mediated gene transfer (Hutson TH et al. 2012; Ueno M et al. 2018) have been used to label axons and terminals of CST neurons in the spinal cord. These studies and earlier tracing studies (Casale EJ et al. 1988) showed that the CST axons terminate mainly in the laminae III and IV of the dorsal horn, where they contact dorsal horn interneurons (Abraira VE et al. 2017; Ueno M et al. 2018). However, the Emx1 or Thy1-H fluorescent reporter mice used in these studies label many neurons in addition to CST neurons (Bareyre FM et al. 2005; Porrero C et al. 2010; Willenberg R and O Steward 2015; Zeisel A et al. 2015) and therefore do not allow specific targeting of CST neurons for characterization and functional manipulation. To this end, it will be crucial to restrict transgene expression to the layer 5 pyramidal neurons in an area of the cortex (S1 in this case) that projects to the spinal region of interest such as the dorsal horn. Recently, a new recombinant adeno-associated virus (rAAV) serotype (rAAV2-retro) (Tervo DG et al. 2016) has been developed with greatly improved retrograde labelling efficacy that allows high fidelity tracing of descending inputs to the spinal cord (Haenraets K et al. 2017; Wang Z et al. 2018).

Here, we developed a combination of viral approaches and transgenic mice to specifically label S1-CST neurons. This approach permitted the expression of fluorescent or effector proteins in S1 cortical neurons that project directly to a predefined region of the spinal cord and allowed us to demonstrate that S1-CST neurons with terminations in the spinal dorsal horn constitute a heterogeneous population of neurons that receive monosynaptic input from forebrain sensory circuits and target dorsal horn interneurons known to be involved in the gating of somatosensory and nociceptive signals.

## Methods

### Animals

Experiments were performed on 6-12-week-old mice kept at a 12:12 h light/dark cycle. Permissions for experiments have been obtained from the Canton of Zurich (permissions 03/2018, 031/2016, and 063/2016). CCK^Cre^ mice (Cck<tm1.1(cre)Zjh>/J, (Taniguchi H et al. 2011)) were purchased from Jackson Laboratory. For further details on the genetically modified mice used in this study, see Table 1.

**Table 1.**
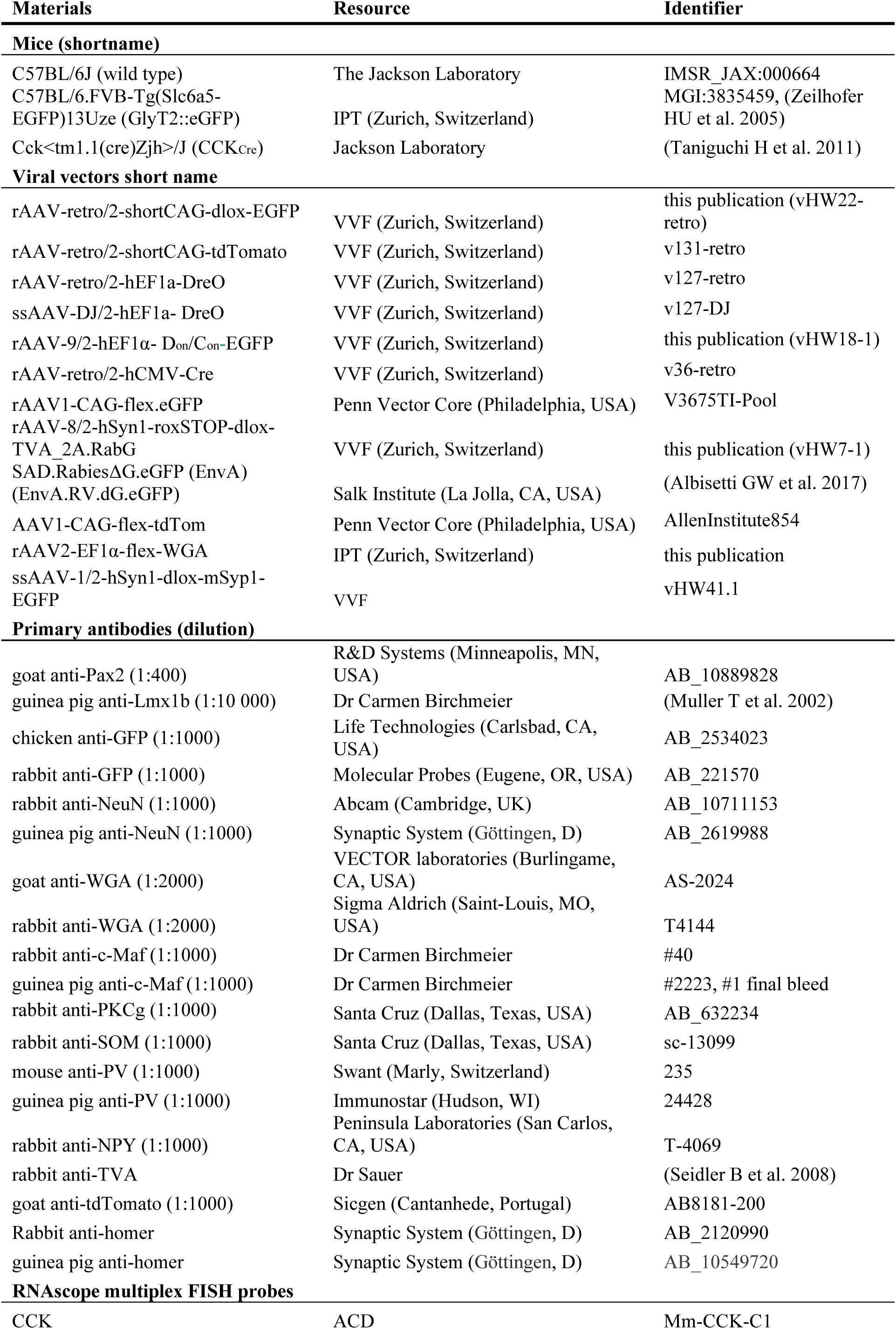
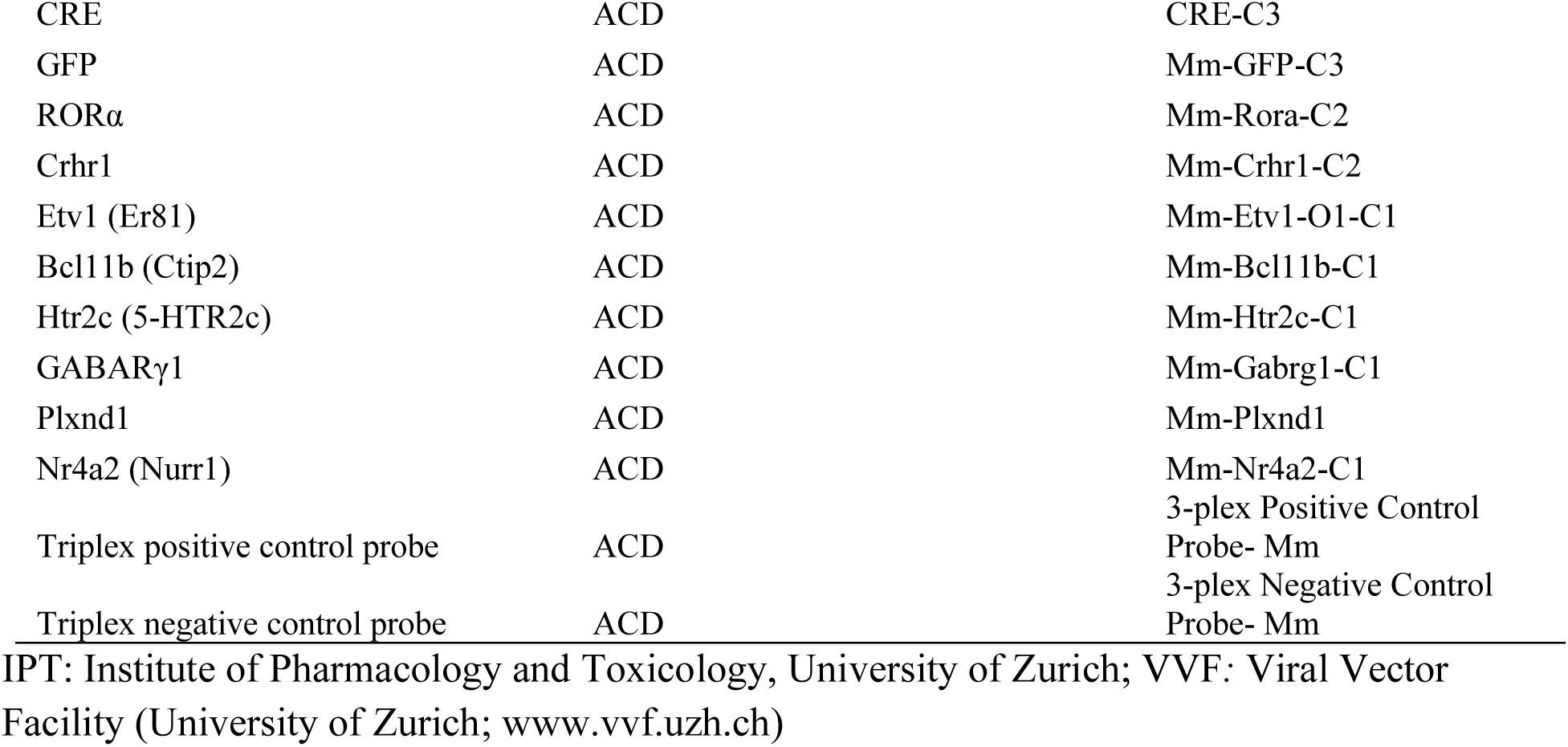
Materials and reagents

### Immunohistochemistry (IHC)

Mice were transcardially perfused with 4% ice-cold paraformaldehyde (in 0.1M sodium phosphate buffer, pH 7.4). Lumbar spinal cord and brain were immediately dissected and post-fixed for 2.5 hrs with 4% paraformaldehyde (PFA) on ice. Post-fixed tissue was briefly washed with 0.1M sodium phosphate buffer (pH 7.4) and then incubated in 30% sucrose (in PBS) overnight at 4°C for cryoprotection. Cryoprotected tissue was cut at 25 μm or 40 μm (spinal cord or brain, respectively) on a Hyrax C60 Cryostat (Zeiss, Oberkochen, Germany), mounted on superfrost plus glass slides and then incubated with the respective combinations of primary antibodies in 1% donkey serum in phosphate buffered saline (PBS) over-night at 4°C. After brief washes in PBS, sections were incubated with the respective secondary antibodies for 2 hrs at room temperature and briefly rinsed in PBS, before mounting with coverslips and DAKO fluorescent mounting media (Dako, Carpinteria, CA, USA). Secondary antibodies raised in donkey were purchased from Jackson Immuno Research (West Grove, PA, USA). All primary antibodies used are listed in the Table 1.

### Multiplex in situ hybridization (ISH) and image analysis

Spinal tissue used for ISH was dissected from 6-12-week-old mice, collected in 1.5 ml Eppendorf tubes, and immediately frozen in liquid nitrogen. Tissue was embedded in NEG50 frozen section medium (Richard-Allen Scientific), cut into 16 μm thick sections, and hybridized using the probes designed for RNAscope Fluorescent Multiplex ISH listed in Table 1.

For IHC and ISH, Z-stacks of fluorescent images were acquired on a Zeiss LSM700 confocal and a Zeiss LSM800 Airy Scan microscope (Zeiss, Oberkochen, Germany). Numbers of immunoreactive cells in z-stacks were determined using the ImageJ (NIH, Bethesda, Maryland) Cell Counter plugin (Kurt DeVos, University of Sheffield, Academic Neurology).

### Intraspinal and cortical virus injections

Viruses were obtained from the resources indicated in Table 1 and used as previously described (Haenraets K et al. 2017). Virus injections were made in adult (6-8-week-old) mice anesthetized with 2% isoflurane and immobilized on a motorized stereotaxic frame (David Kopf Instruments, Tujunga, CA, USA and Neurostar, Tübingen, Germany). For intraspinal injections, the vertebral column was fixed using a pair of spinal adaptors and lumbar spinal cord at L4 and L5 was exposed. Injections (3 × 300 nL) spaced approximately 1 mm apart were made at a rate of 50 nl/min with glass micropipettes (tip diameter 30 - 40 μm) attached to a 10 μl Hamilton syringe. For S1 injections, the head was fixed using head bars, the skull exposed and the following injection coordinates were used: (Bregma -1 mm; midline +1.5 mm; depth: 0.8 mm).

### Tissue clearing and light sheet imaging

In order to label the CST neurons more sparsely, we injected an AAV-DJ/2.Dre into the lumbar spinal cord instead of the rAAV2-retro in order to get sparse labelling of CST neurons and better visualisation of individual neurons and axons in cleared brains. Three weeks after viral injections, mice were deeply anesthetized using pentobarbital and perfused transcardially with 10 ml of ice cold PBS followed by 20 ml of ice cold hydrogel solution (40% acrylamide, 2% bis-acrylamide, 10% VA-044 initiator, PBS, 16% PFA and dH_2_O (Chung K et al. 2013). Whole CNS (brains and with spinal cords attached) were dissected on ice and incubated in hydrogel solution for 24 hrs. Oxygen was removed using a desiccator and N_2_ used to replace it to allow a good polymerization. Sampled were incubated for 3 hrs at 39.5°C for acrylamide polymerization, and then incubated in SDS (8% SDS, 200 mM boric acid and sodium hydroxide to adjust the pH at 8.5 (Chung K et al. 2013), at RT for passive clearing. After 8-9 weeks, the samples were washed in PBS at least 3 times for 4 hrs and then incubated in Histodenz-Triethanolamine solution (about 80% Histodenz D2158 (Sigma-Aldrich), 11.5 mM PB, 5.7% Na-azide, 5.7% Tween 20 and ∼30% Triethanolamine, pH 7.5; refractive index adjusted to 1.457) for 7 days. Finally, samples were mounted in quartz cuvettes (10 × 20 × 45 mm^3^, Portmann Instruments) for imaging in the same solution. Images were acquired using a mesoSPIM light-sheet microscope (‘mesoscale selective plane illumination microscope’, see mesospim.org) (Voigt FF et al. 2019). Briefly, samples were illuminated with an axially scanned light-sheet using an Omicron SOLE-6 laser engine at 488-nm and 561-nm excitation wavelengths. 3D stacks were generated by translating the sample through the light sheet. In the detection path, fluorescence signals were acquired with an Olympus MVX-10 microscope with a MVPLAPO 1x objective and a Hamamatsu Orca Flash 4.0 CMOS camera. Image analysis was performed with Fiji and Imaris (version 9.5.1, Bitplane AG).

### Quantification of retrogradely labelled neurons

For the quantification of the retrogradely labelled neurons using rabies virus, percentage of PV and SST neurons were counted as percentage of GFP^+^TVA^-^ neurons. Because of the lack of suitable antibodies, it was not possible to co-stain for NPY and TVA in the same section. Therefore percentage is expressed as percentage of all GFP^+^ neurons. However, since we only found a single NPY and GFP positive neuron in the brains of 3 mice (196 neurons), we are confident that the main conclusions are not affected. Finally, pyramidal layer 2/3 neurons were identified based on their localization and morphology.

### Experimental design and statistical analysis

Cells counts are reported as mean ± SEM. Numbers of experiments (mice and cells) are provided in the figure legends.

## Results

### Labelling S1-CST neurons in CCK^Cre^ mice

CST neurons are the subset of excitatory long-range pyramidal projection neurons in layer 5 of the somatosensory cortex that terminate in the dorsal spinal horn where they release glutamate as their fast neurotransmitter. However, whether S1-CST neurons constitute a molecularly or functionally homogeneous population is unclear. Recent work (Zeisel A et al. 2015) indicates that the neuropeptide cholecystokinin (CCK) is also expressed in some layer 5 neurons of S1. We therefore addressed the question whether and, if so, to what extend CST neurons in S1 express CCK. To specifically label CCK expressing CST neurons in S1, we decided to inject transgenic mice carrying a knock-in Cre coding sequence in the CCK locus (CCK^Cre^ mice, (Taniguchi H et al. 2011)) with a rAAV optimized for transduction through axons and axon terminals (rAAV2-retro). We have previously shown that spinal injection of rAAV2-retro vectors into the lumbar spinal dorsal horn transduces CST neurons in the primary sensory cortex (S1) via infection of their spinal axon terminals (Haenraets K et al. 2017). We therefore repeated our initial experiments in CCK^Cre^ mice. First, we confirmed that Cre expression in these mice is restricted to CCK expressing cells (98.8 ± 0.4% of Cre^+^ cells expressed CCK and 98.6 ± 0.7% of CCK^+^ cells expressed Cre, Fig. S1) using multiplex in situ hybridization. We then injected a rAAV2-retro encoding a Cre-dependent eGFP (AAV2-retro.flex.eGFP) into the spinal cord of these mice (Fig. 1A). This strategy uncovered a population of layer 5 pyramidal neurons in the primary somatosensory cortex that projects directly to the spinal dorsal horn (Fig. 1B). We also found labelled neurons in a few other brain areas, including the anterior cingulate cortex, the thalamus, and the rostral ventromedial medulla (RVM) (Fig. 1B). However, in all subsequent experiments, we focused on our initial central aim, the characterization of CST neurons in the somatosensory cortex. We tested whether all CST neurons in S1 expressed CCK^Cre^ or whether the CCK^Cre^ neurons were a subset of CST neurons. To this end, we co-injected a Cre dependent (rAAV2-retro.flex.eGFP) and a Cre independent rAAV2-retro (rAAV2-retro.tdTomato) into the lumbar spinal cord of CCK^Cre^ mice (Fig. 1C). eGFP would then be expressed in CCK^Cre^ positive S1-CST neurons, whereas tdTom would label all virus infected neurons projecting to the injection site. We found that about 70% (71.5 ± 3.1%) of the tdTom^+^ S1-CST neurons also expressed eGFP. The proportion eGFP^+^ neurons that expressed tdTomato was very similar (75.5 ± 1.6%, Fig. 1D). These values are consistent with an expression of CCK^Cre^ in the great majority of S1-CST neurons that project to the injection site. S1-CST neurons can hence be efficiently labelled using rAAV2-retro injections into the dorsal horn of CCK^Cre^ mice. CCK is not only expressed in cortical layer 5 pyramidal neurons of the mouse cortex but also in large subsets of excitatory and inhibitory neurons of the cerebral cortex (Xu X et al. 2010; Zeisel A et al. 2015). This also became apparent when we injected rAAVs containing a Cre-dependent tdTomato expression cassette directly into the S1 cortex (Fig. 1E-F). As a consequence, injection of rAAVs carrying Cre dependent reporter cassettes into CCK^Cre^ mice allows anterograde labelling of S1-CST terminals from the cortex and retrograde labelling of their somata from the spinal cord (Fig. 2A). It does however not permit a specific transduction of CST neurons from either of the two injection sites alone.

**Fig. 1:**
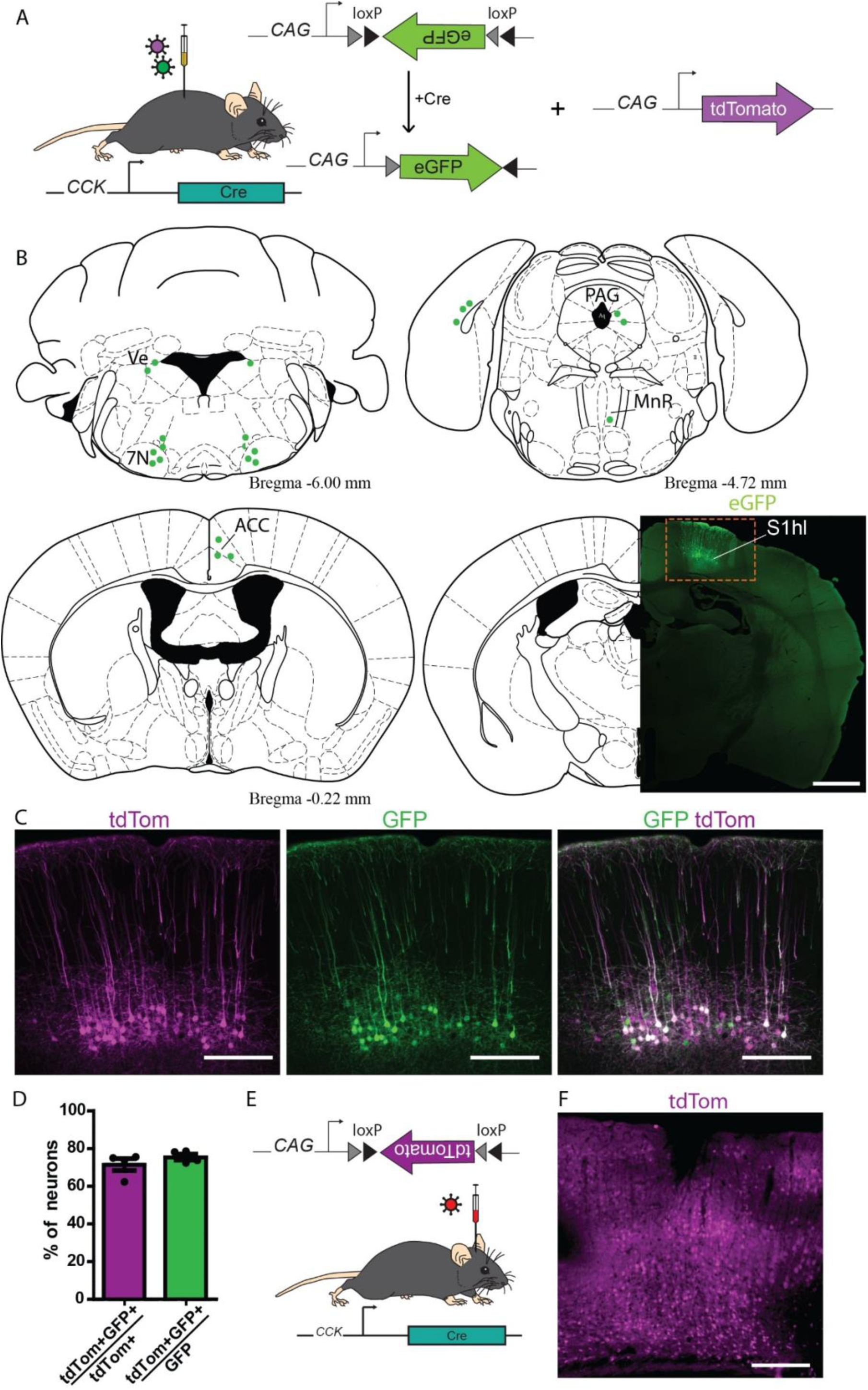
Labelling S1-CST neurons in CCK^Cre^ mice. (A) Injection of rAAVs encoding for Cre-dependent eGFP and Cre-independent tdTomato fluorescent proteins into the lumbar spinal cord of CCK^Cre^ mice (n = 4 mice). (B) Brain areas labelled with eGFP positive neurons after intraspinal injection of rAAV2retro.flex.eGFP in CCK^Cre^ mice. 7N: facial nuclei, ACC: anterior cingulate cortex, MnR: median raphe nucleus, PAG: Periaqueductal grey, S1hl: somatosensory cortex, hindlimb area. (C) Comparison of S1-CST neurons labelled by Cre-dependent eGFP and Cre-independent tdTomato (n = 4, 3166 neurons) fluorescent proteins. (D) Quantification of (C). (E) Injection of rAAVs encoding for Cre-dependent tdTomato into the S1 cortex of CCK^Cre^ mice. (F) Widespread labelling of cortical neurons with tdTomato (red) after cortical injection (E). Error bars represent ± SEM. Scale bars: B: 1 mm; C and F: 200 µm.

**Fig. 2:**
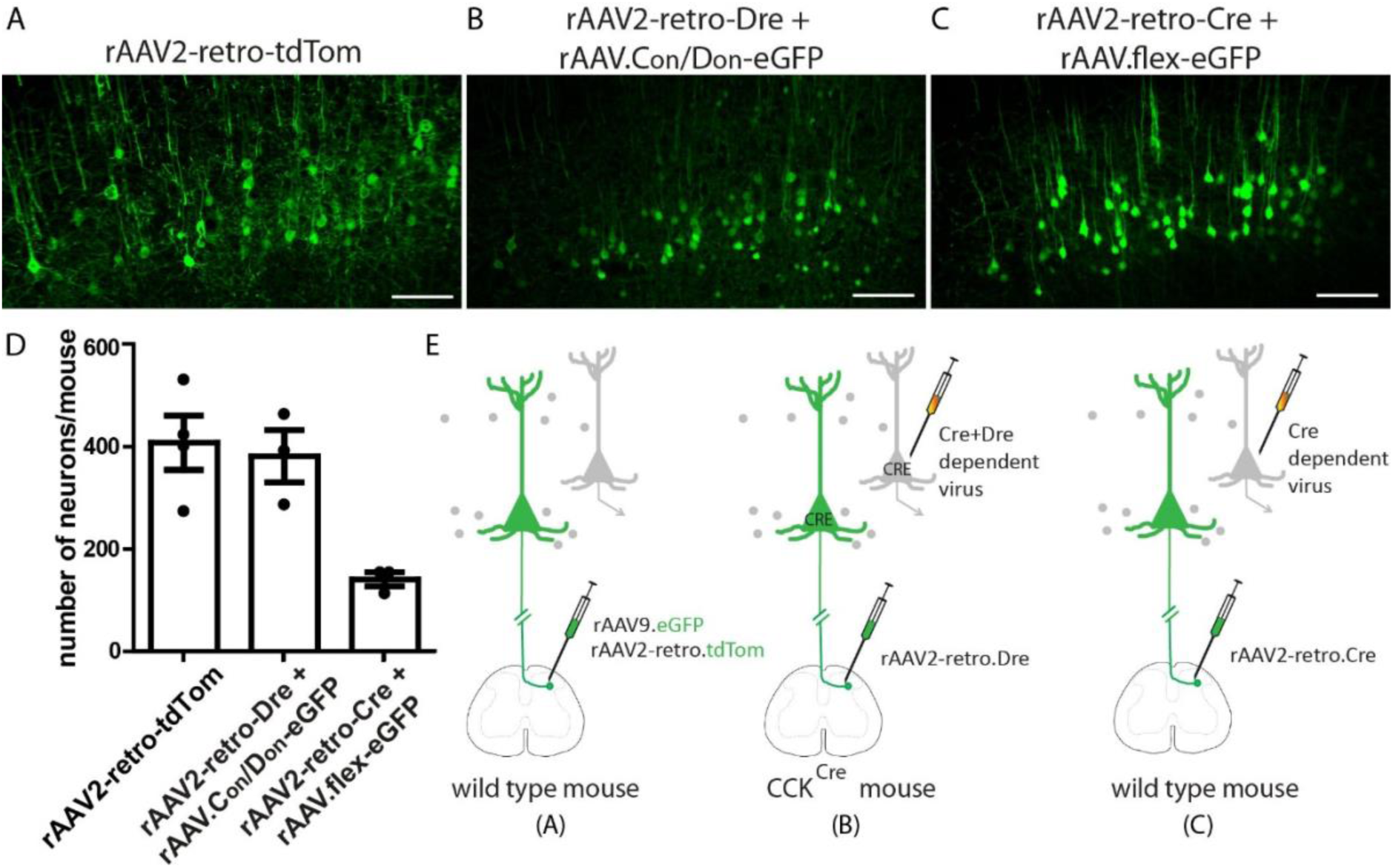
Three viral targeting strategies to label S1-CST neurons. (A) S1-CST neurons labelling using a rAAV2-retro-tdTomato injected in the lumbar spinal cord of wild-type mice. (B) S1-CST neurons labelling using the rAAV2-retro-Dre injected in the lumbar spinal cord of CCK^Cre^ mice, followed by cortical (S1) injection of AAV.C_on_/D_on_-eGFP. (C) S1-CST neurons labelling using the rAAV2-retro-Cre injected in the lumbar spinal cord of wild-type mice, followed by cortical (S1) injection of AAV.flex-eGFP. (D) Quantification of the number of neurons labelled per mouse in (A-C) (A: n = 4 mice, 1546 neurons; B: n = 3 mice, 1136 neurons and C: n = 3 mice, 418 neurons). (E) Diagrams showing the three injections strategies used in (A), (B) and (C), respectively. Error bars represent: ± SEM. Scale bars: 100 µm.

### Viral targeting strategies to label S1-CST neurons

To overcome this limitation, we developed an intersectional strategy (Fig. 2) to specifically target CCK^+^ neurons that project from S1 to the spinal cord. To this end, we injected a rAAV2-retro encoding for the Dre recombinase into the lumbar spinal cord of CCK^Cre^ mice (Fig. 2B). Subsequently, transduced S1-CST neurons (as well as other CCK^Cre^ neurons with processes or somata in the spinal cord) will express both recombinases. Using this strategy, we achieved targeted expression of the desired transgene by local injection of rAAVs carrying Cre-and Dre-dependent transgenes into S1. As a proof of principle, we demonstrate that this strategy works with the injection of a Cre- and Dre-dependent rAAV carrying the eGFP transgene (AAV.hEF1α.C_on_/D_on_.eGFP) into S1 (Fig. 2B). We did not detect eGFP expression outside of S1.

A possible variation of this intersectional strategy is the injection of a rAAV2-retro.Cre into the spinal cord of wild-type mice, followed by the local injection of a rAAV carrying a Cre-dependent transgene into S1 (Fig. 2C). As CCK^Cre^ positive neurons represent the vast majority of the S1-CST population, the two strategies should label the same neuron population in this particular case. Quantification of the different injection strategies shows that a single injection of rAAV2-retro.tdTom into the spinal cord of wild type mice (Fig. 2A, D) led to labelling of a similar number of neurons as compared to the C_on_/D_on_ strategy in CCK^Cre^ mice (Fig. 2B, D). Targeting S1-CST neurons by spinal injection of rAAV2-retro.Cre in wild-type mice, followed by the local injection of a rAAV.flex-eGFP (Fig. 2C) led to labelling of less than half as many neurons (Fig. 2D).

### Molecular characterization of S1-CST neurons

Since virtually all S1-CST neurons projecting to the lumbar spinal cord express CCK, we next asked whether these neurons constitute a homogeneous population or are assembled from different subpopulations. To this end, we injected a rAAV2-retro.flex.eGFP into the spinal cord of CCK^Cre^ mice and performed multiplex *in situ* hybridization against several established markers for cortical neurons in cortex sections obtained from the virus-injected mice (Fig. 3A-E). Consistent with our initial characterization of the CCK^Cre^ mice, we found that all eGFP-labelled neurons expressed the CCK mRNA (Fig. 3A). Furthermore, the vast majority of eGFP labelled CST-neurons expressed well-established marker genes of cortical layer 5 neurons (Arlotta P et al. 2005; Molyneaux BJ et al. 2007; Watakabe A et al. 2007; Klingler E et al. 2019) including *Crhr1* (78.9 ± 4.9%), Er81 (79.2 ± 6.5%), or *Ctip2* (80.8 ± 2.5%) (Fig. 3A, 3B, 3C, respectively). We also found that 48.2 ± 4.0% of eGFP^+^ neurons expressed *RORα* and 23.5 ± 1.4% expressed *Nurr1* (Fig. 3D) mRNAs, indicating molecular heterogeneity within the CCK^+^ CST neurons. Expression of several other genes was detected at low levels and only in few eGFP^+^ neurons (*gabrg1*: 12.8 ± 2.3 %, *5HT2c*: 10.37 ± 3.6 %, and *Plxnd1*: 4.33 ± 1.4 %, Fig. 3D and 3E, RNAscope negative probe background level is depicted in Fig. S2).

**Fig. 3:**
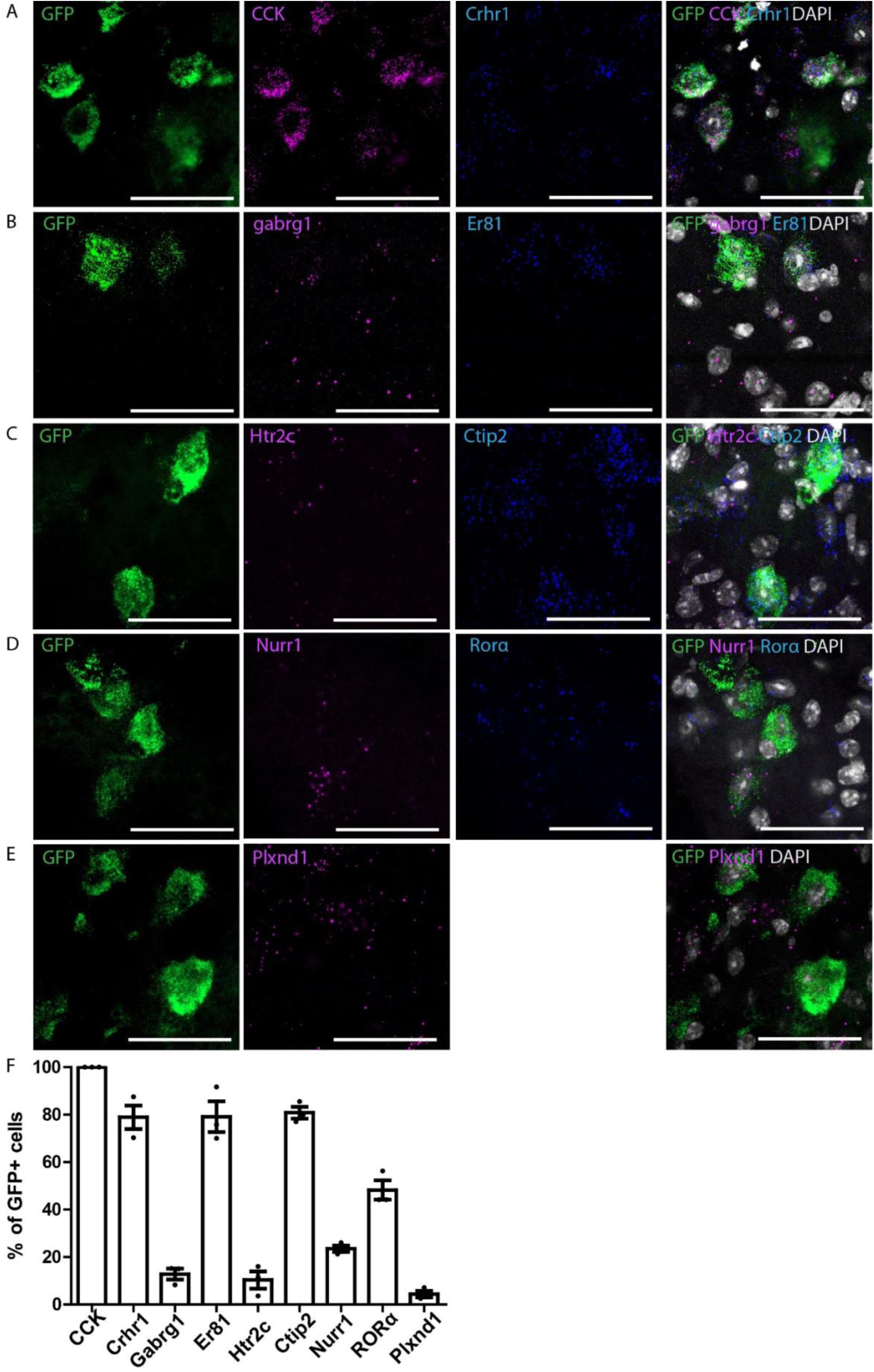
Multiplex ISH in GFP-labelled CCK^Cre^ neurons with cortical neurons markers in S1. (A-E) Triple ISH showing the colocalization of GFP with *CCK* and *crhr1* (A), *gabrg1* and *er81* (B), *htrc2* and *ctip2* (C), nurr1 and *RORα* (D), *Plxnd1* (E). (F). Quantification of (A-E) (n = 3 mice; 352, 221, 85, 243, and 278 GFP neurons respectively). Error bars represent ± SEM. Scale bars: 50 µm.

### Morphology of S1-CST neurons

We next examined whether S1-CST neurons send collaterals to other central nervous system (CNS) regions before they reach the spinal cord. We employed the intersectional strategy described above to label CCK^Cre^ S1-CST neurons (Fig. 2B). Three weeks after the virus injections, entire mouse CNS were dissected and cleared using the passive CLARITY procedure (Chung K et al. 2013). Cleared entire mouse CNS were imaged with light-sheet microscopy (Voigt FF et al. 2019). The vast majority of S1-CST neuron axons ran from the cortex through the internal capsule and to the midbrain pyramids, following the known trajectory of the main CST (Fig. 4) (Wang X et al. 2017). In addition, we detected a few collaterals branching from the main tract at two sites: a small number of axons bifurcated from the internal capsule to terminate in the dorsal striatum (Fig. 4B, C), and another small group branched-off in the midbrain innervating posterior thalamic nuclei (Fig. 4D) and tectal areas (Fig. S3D-E). The latter is consistent with observations from Wang et. al in the motor cortex-derived CST (Wang X et al. 2017). As expected, the CST axons travel from a ventral to a dorsal location as they leave the brainstem to enter the spinal cord. At this level the tract is also moving to the contralateral side of the spinal cord (pyramidal decussation, Fig. 4E). We observed a few axons branching into the dorsolateral CST (Fig. 4E-G, I-J) (Steward O et al. 2004; Kathe C et al. 2014). In the lumbar spinal cord, axons bifurcate from the CST into the dorsal horn grey matter (corresponding to the location of their spinal targets and the site of injection of the Dre carrying virus). We did not observe collaterals into segments of the spinal cord other than the lumbar segments (Fig. 4H, K).

**Fig. 4:**
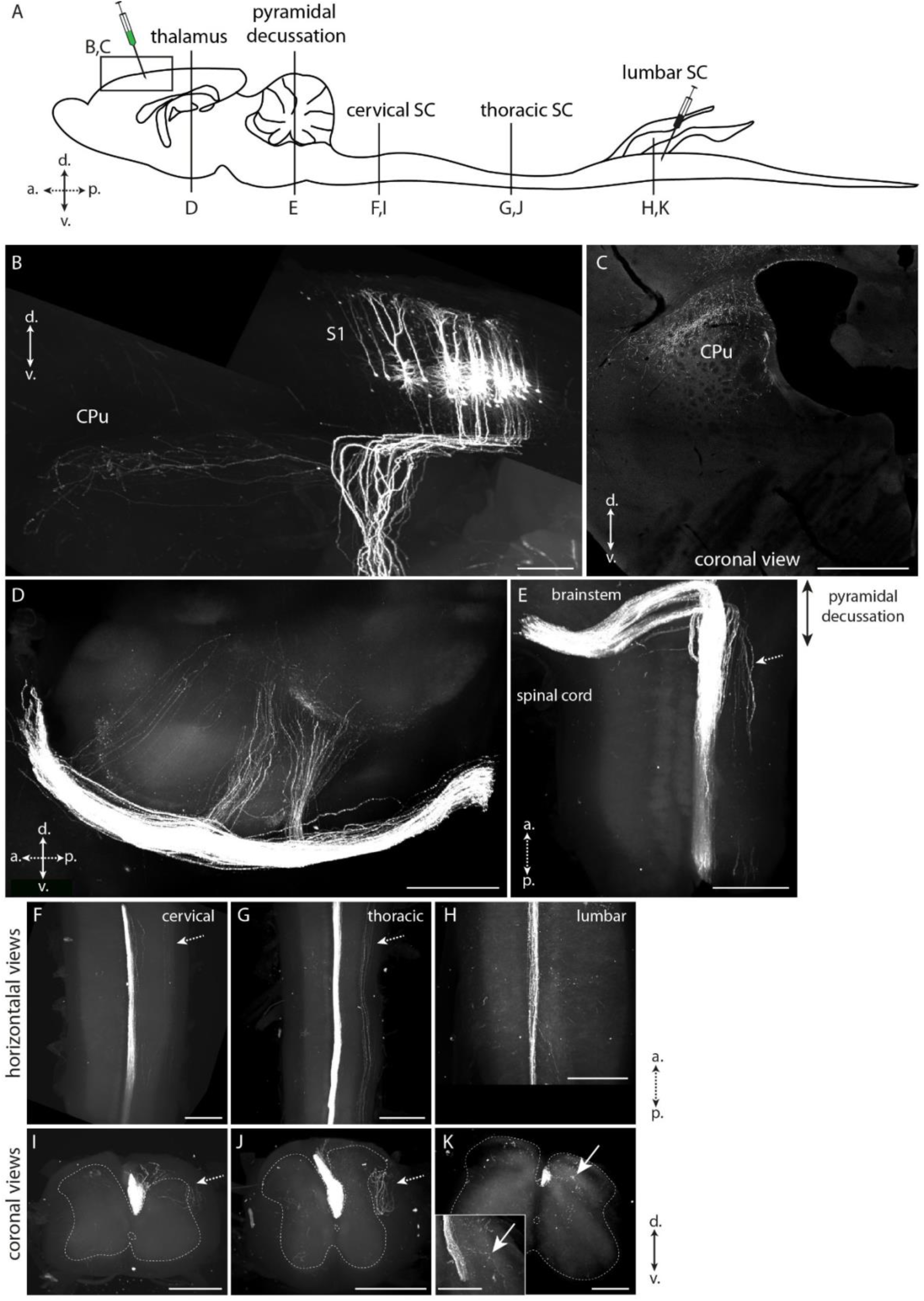
Labelling of the corticospinal tract in CLARITY-cleared brain. The whole CNS of mice expressing eGFP only in lumbar spinal cord projecting CST neurons was dissected and subjected to passive clearing and light sheet imaging. (A) Schematic drawing of the mouse CNS. Injection sites of the viruses and optical planes shown in B-K are depicted. (B) eGFP labelling of S1-CST neurons in S1. The axons enter the corpus callosum and a small subset of collaterals bifurcates to the dorsal striatum (CPu). (C) Coronal view of the termination area in the CPu. (D) Sagittal view of collaterals branching of from the main CST towards the thalamus. (E) Horizontal view of the CST decussation at the entry of the spinal cord. (F, I) Horizontal and coronal views of the CST at the level of the cervical spinal cord. (G, J) Horizontal and coronal views of the CST at the level of the thoracic spinal cord. (H, K) Horizontal and coronal views of the CST at the level of the lumbar spinal cord. Inset in (K) shows CST terminals branching in the dorsal horn. White arrows: CST terminals branching in the spinal cord at the lumbar level only. Dashed arrows: dorsolateral tract (secondary part of the CST). CPu: Striatum; S1: primary somatosensory cortex (hindlimb area here); SC: spinal cord; a.: anterior, p.: posterior, d.: dorsal, v.: ventral. B, D-K: Volume renderings (Imaris); C: optical section. (n = 3 mice). Scale bars: 1 mm.

### S1-CST neurons receive input from the somatosensory circuit

The direct connection between the spinal dorsal horn and the somatosensory cortex suggests that S1-CST neurons may be part of a circuit for sensory processing. We therefore further investigated the precise position of S1-CST neurons in this circuit by tracing their presynaptic input and postsynaptic target neurons. We started with the identification of neurons presynaptic to S1-CST neurons and performed monosynaptic retrograde labelling using genetically engineered rabies virus (Callaway EM and L Luo 2015). In order to label as many neurons as possible (Fig. 2D), S1-CST neurons were targeted in CCK^Cre^ mice by intraspinal injection of rAAV2-retro.Dre followed by local injection of a Cre and Dre dependent helper virus (rAAV.flex.rox.TVA.SAD19-G) into the primary somatosensory cortex S1 (Fig. 5). The helper virus provided the TVA gene for selective infection by EnvA-pseudotyped rabies virus and the rabies glycoprotein (SAD19-G) for trans-complementation enabling retrograde labelling across one synapse. We then injected a glycoprotein-deficient EnvA-pseudotyped rabies virus (EnvA.RV.ΔG.eGFP) into the S1 cortex. We found that eGFP was expressed in layer 5 pyramidal neurons (including the primarily infected S1-CST neurons, Fig. 5A) and in many pyramidal neurons of layer 2/3 (Fig. 5E-F). In layer 5, the rabies virus also labelled interneurons that expressed parvalbumin (PV, 15.4 ± 2.73% of GFP^+^TVA^-^ neurons), and rarely somatostatin (SOM; 0.77 ± 0.41%) or NPY (0.74 ± 0.74%) (Fig. 5G-I), three well-characterized markers of cortical inhibitory interneurons (Xu X et al. 2010), representing approximatively 60-70% of cortical inhibitory interneurons (Gonchar Y et al. 2007).

**Fig. 5:**
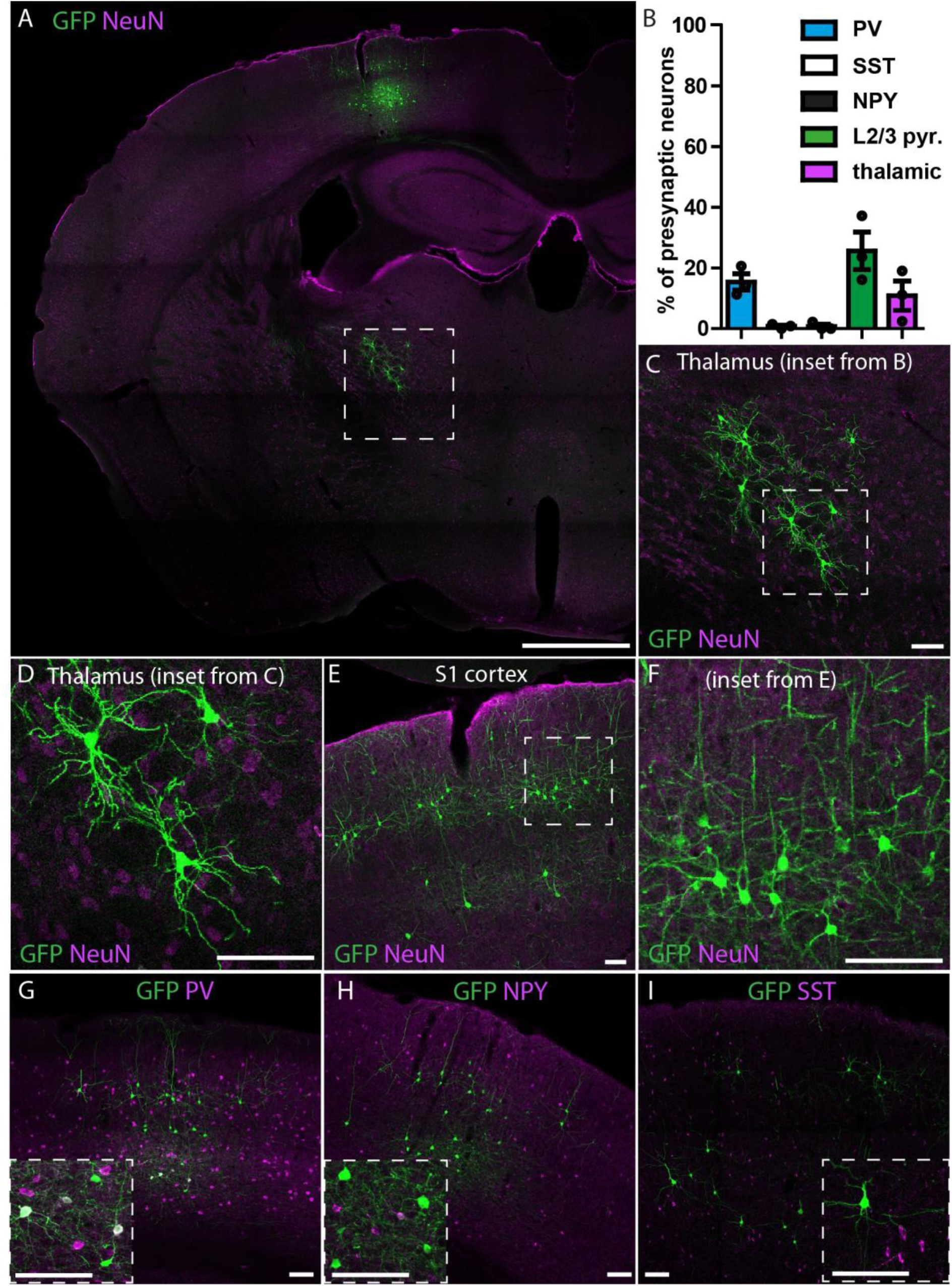
Retrograde monosynaptic tracing of S1-CST neurons with rabies. rAAV2retro.Dre was injected into the spinal cord of CCK^Cre^ mice, followed by a Cre-and-Dre-dependent helper virus (TVA, RabG) into S1. Two weeks later the pseudotyped rabies virus was injected into S1. (A) Overview of the labelled neurons in the brain: S1-CST neurons (starter cells) as well as layer 2/3 pyramidal neurons, thalamic sensory relay neurons and layer 5 inhibitory interneurons. (B) Quantification of retrogradely labelled neurons (GFP^+^TVA^-^) represented in (C-I). (C-D) Retrogradely labelled thalamic sensory relay neurons in the VPL of the thalamus (n = 3 mice, 1481 GFP^+^ neurons). (E-F) Retrogradely labelled layer 2/3 pyramidal neurons (n = 3 mice, 1481 GFP^+^ neurons). (G-I) Retrogradely labelled layer 5 inhibitory interneurons, with co-staining against PV (G, n = 3 mice, 513 neurons), NPY (H, n = 3 mice, 196 GFP^+^ neurons) or SST (I, n = 3 mice, 968 GFP^+^ neurons). Error bars represent ± SEM. Scale bars: A: 1 mm; C-I: 100 µm.

We also found eGFP^+^ neurons in the ventral posterolateral nucleus (VPL) of the thalamus (Fig. 5C-D, 10.9 ± 4.8% of GFP^+^TVA^-^ neurons). The morphology of these cells resembled that of previously described thalamocortical sensory relay neurons (Tlamsa AP and JC Brumberg 2010). This connectivity pattern is hence similar to what has previously been described for CST neurons in other studies (Abraira VE and DD Ginty 2013; Constantinople CM and RM Bruno 2013; McMahon et al. 2013). Our results thus demonstrated that CCK-expressing S1-CST neurons are part of a direct sensory circuit loop between the spinal cord, thalamic nuclei and the somatosensory cortex.

### Labelling of CST axons in the spinal cord

The termination area of S1-CST neurons in the lumbar spinal cord (Fig. 6) was studied after injection of rAAV1.flex.tdTomato in the somatosensory cortex hindlimb area (S1hl) of CCK^Cre^ mice. Labelled CST axons were mainly found in the ventral part of the spinal dorsal funiculus (Fig. 6A, “CST”). Terminals were also visible within the grey matter of the deep dorsal horn (laminae III and IV) (Fig. 6A, S4B-C). This finding is consistent with previous reports showing that tracing from the motor cortex labeled terminals mainly in the ventral and intermediate spinal cord, whereas tracing from S1 labeled terminals in lamina III and IV of the dorsal horn (Kamiyama T et al. 2015; Wang X et al. 2017; Ueno M et al. 2018). These dorsal horn laminae constitute the termination area of low-threshold mechanoreceptive fibres (LTMRs) (Abraira VE et al. 2017). In addition, they contain interneurons which process touch and proprioceptive information, and have been shown to be critically involved in abnormal somatosensory processing in neuropathic pain (Foster E et al. 2015; Peirs C et al. 2015; Petitjean H et al. 2015; Cheng L et al. 2017). We therefore decided to identify the spinal neurons that are targeted by S1-CST neurons in this region.

**Fig. 6:**
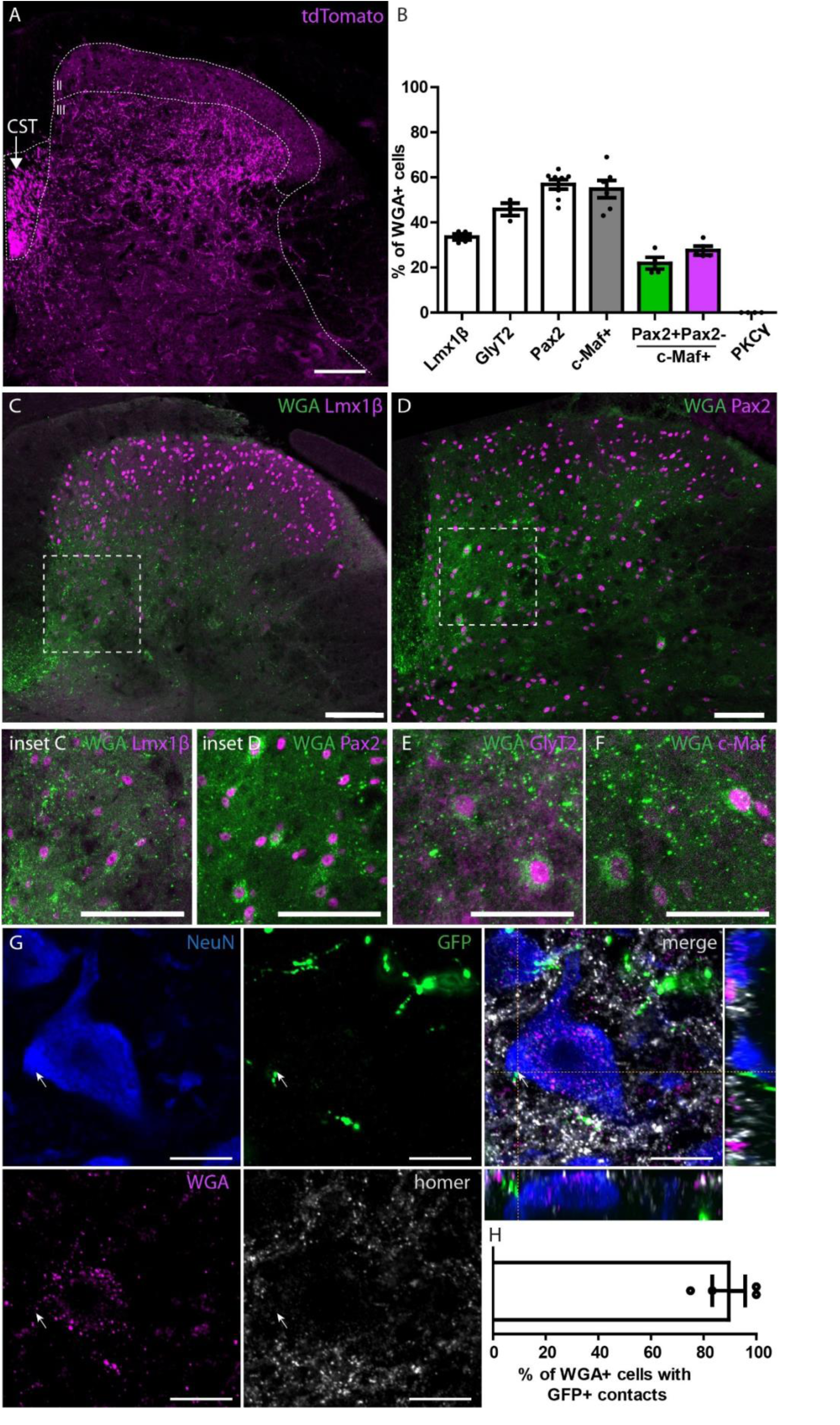
Labelling of the output of S1-CST neurons in the spinal cord. (A) Labelling of the CST in the dorsal funiculus of the spinal cord, contralateral to the brain injection site after injection of a rAAV carrying a Cre-dependent tdTomato into S1hl of CCK^Cre^ mice. CST terminals are preferentially located below the laminae II-III border (n = 3). (B) Quantification of the number of WGA positive neurons after injection of a rAAV.flex.WGA into S1hl of CCK^Cre^ mice. Quantified are WGA positive neurons that express Lmx1β (n = 4, 320 WGA^+^ neurons), Pax2 (n = 8, 391 WGA^+^ neurons), GlyT2 (n = 3, 275 WGA^+^ neurons), c-Maf (n = 6, 506 WGA^+^ neurons) or PKCγ (n = 4, 201 WGA^+^ neurons). (C-D) Representative images of co-labelled WGA positive neurons in the spinal cord with the excitatory marker Lmx1β ((C) and inset) and the inhibitory marker Pax2 ((D) and inset). Neurons expressing eGFP under the GlyT2 promoter (using the GlyT2::eGFP mouse line, (E)), and neurons expressing the transcription factor c-Maf (F) were also found positive for WGA. (G-H) Verification of monosynaptic labelling by WGA. CCK^Cre^ Mice were co-injected with rAAV.flex.WGA and rAAV.flex.Syp-eGFP (encoding a Cre-dependant synaptophysin-eGFP fusion protein) into S1hl. (G) Co-labelling of WGA positive neurons in the spinal cord with the neuronal marker NeuN and eGFP labelled presynaptic terminals of S1-CST neurons. Depicted is a representative example of a WGA^+^ neuron in close proximity to a eGFP^+^ presynaptic terminal of a S1-CST neuron3. (H) Quantification of the number of WGA^+^ neurons receiving direct contacts from eGFP^+^ synaptic terminals (n = 4 mice; 25 neurons). CST: corticospinal tract. Error bars represent ± SEM, Scale bars: A and C-F: 100 µm; G: 10 µm.

### Anterograde transsynaptic tracing with Wheat Germ Agglutinin (WGA)

We used WGA to label neurons that are targeted by S1-CST neurons. CCK^Cre^ mice were injected with rAAV2.flex.WGA into S1hl. WGA is transported transsynaptically to label postsynaptic neurons (Braz JM et al. 2002). After 10 days, we detected WGA in the dorsal horn of the lumbar spinal cord (Fig. 6C-F). As expected, WGA immunoreactivity was mostly found in the deep dorsal horn laminae III and IV (Fig. 6C-F, S4B-C). In order to identify the neurons labelled with WGA, we stained sections with antibodies against known markers of different dorsal horn interneuron populations. We found that about one third of the WGA labelled neurons were positive for Lmx1β (33.5 ± 1.4%), which is expressed by most excitatory interneurons in laminae I-III (Del Barrio MG et al. 2013; Albisetti GW et al. 2019) (Fig. 6B-C). More than half of the labelled neurons (56.9 ± 2.1%) were positive for Pax2, a marker of dorsal horn inhibitory neurons (Del Barrio MG et al. 2013; Albisetti GW et al. 2019) (Fig. 6B, D). When performing anterograde tracing in animals crossed to GlyT2::eGFP mice, a little less than half of all WGA positive dorsal horn neurons (45.8 ± 2.8%, Fig. 6B, E) were eGFP^+^ indicating that they were glycinergic (Jursky F and N Nelson 1995; Poyatos I et al. 1997; Spike RC et al. 1997; Zeilhofer HU et al. 2005). Notably, we found that more than half of all WGA positive neurons also expressed the transcription factor c-Maf (54.8 ± 3.9%; Fig. 6B, F), which is required for the proper development of laminae III/IV interneurons (Hu J et al. 2012). Because c-Maf is present in both excitatory and inhibitory dorsal horn interneurons (Hu J et al. 2012; Del Barrio MG et al. 2013), we determined the portion of WGA and c-Maf double-positive neurons that were either inhibitory (Pax2 positive: 21.9 ± 2.6% of all WGA^+^ neurons) or excitatory (Pax2 negative: 27.5 ± 1.9% of all WGA^+^ neurons). We did not find any WGA positive neurons that were also positive for protein kinase Cγ (PKCγ), a marker of a small subpopulation of excitatory dorsal horn neurons located at the border between lamina II and III (Malmberg AB et al. 1997; Polgar E et al. 1999). WGA is an anterograde tracer that can cross multiple synapses. We have limited the incubation time after injection of the rAAV2.flex.WGA to obtain mostly monosynaptic labelling. However, to verify that the majority of WGA labelled spinal neurons received direct inputs from CST neurons, we co-injected viruses carrying a transgene for WGA (rAAV2.flex.WGA) together with a rAAV carrying a transgene encoding a synaptophysin-eGFP fusion-protein into the S1hl. We found that 89.6 ± 6.3% of WGA^+^ neurons showed at least one GFP^+^ apposition close to a homer labelled post-synapse on the cell body (Fig. 6G-H and S2D-E) demonstrating that the vast majority of WGA labelled spinal neurons receive monosynaptic input from CST neurons in S1. In agreement with the absence of WGA labelled PKCγ neurons, we found GFP^+^ contacts on only 6.37 ± 2.7% of PKCγ positive neurons (Fig. S4F). Our results therefore suggest that S1-CST neurons contact a rather heterogeneous population of interneurons in the dorsal horn, including glycinergic neurons and c-Maf expressing neurons.

## Discussion

In the present study, we developed intersectional rAAV-based strategies to characterize S1-CST neurons that innervate the spinal cord. We found that these neurons constitute a genetically heterogeneous group of neurons that in turn innervate a heterogeneous target population in the spinal dorsal horn. Furthermore, we show that they receive direct input from sensory thalamic relay neurons in addition to intracortical synaptic input from layer 2/3 pyramidal and from PV positive interneurons. These results establish a long-range feedback loop between the sensory spinal cord and the output neurons of the primary somatosensory cortex (Fig. 7).

**Fig. 7:**
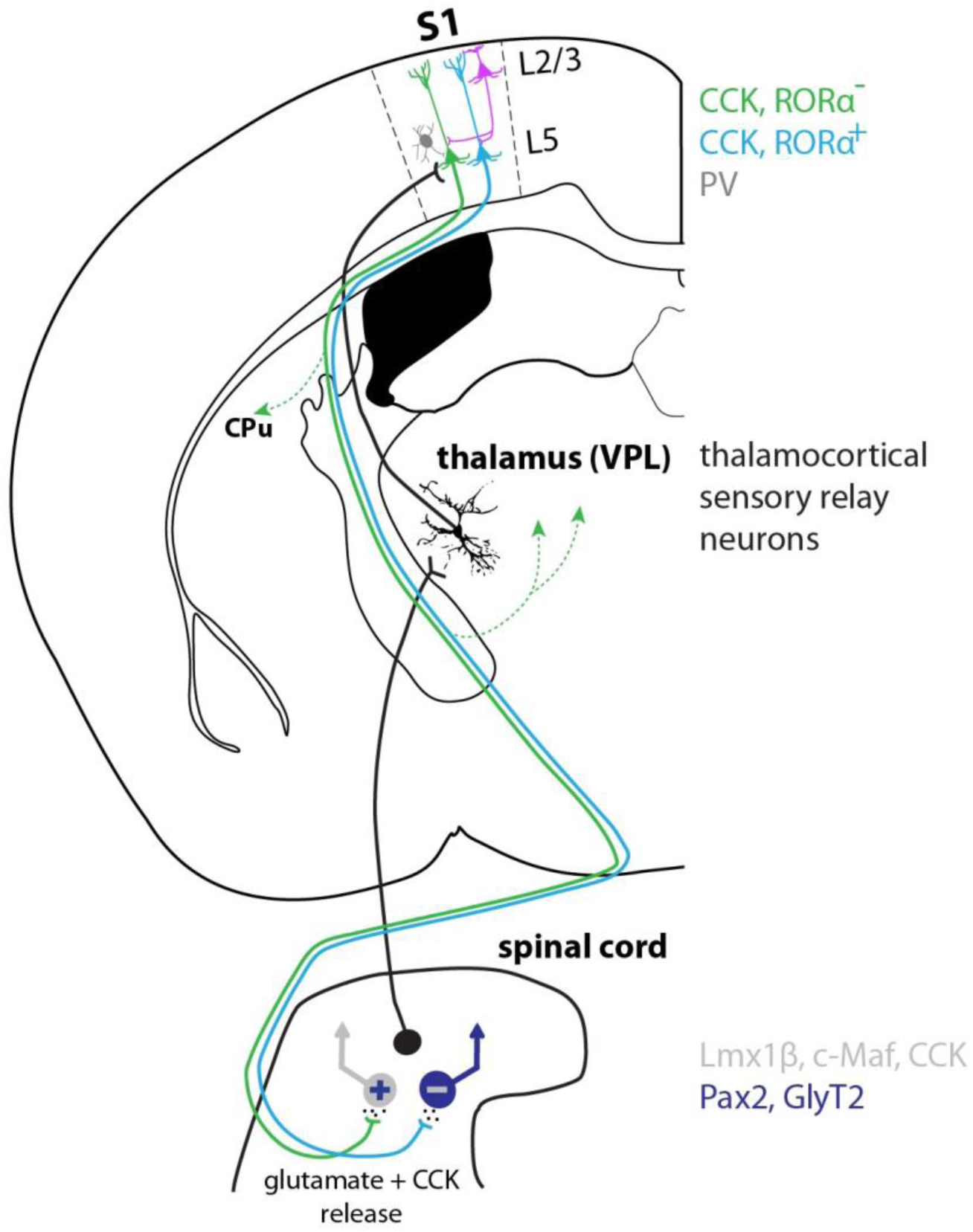
Model of a spinothalamocortical feedback circuit. Spinal projection neurons relay sensory information to the ventral posterolateral nucleus of the thalamus (VPL). Sensory thalamocortical relay neurons propagate the information directly to CST neurons in S1. S1-CST neurons also receive direct synaptic input from inhibitory (PV) neurons and pyramidal layer 2/3 neurons. S1-CST neurons do not only innervate the spinal cord but also send collaterals to the dorsal striatum (CPu) as well as thalamic nuclei (indicated by dotted green lines with arrow). We speculate that different subpopulations of CST neurons (e.g. RORα^+^ or RORα^−^) project back onto different types of spinal interneurons (e.g. inhibitory (GlyT2^+^, Pax2^+^) or excitatory (Lmx1β^+^, CCK^+^, c-Maf^+^) and thus exert potentially modality specific functions (see discussion). CST neurons likely modulate spinal target neurons through release of glutamate as well as the neuropeptide CCK.

### CCK expression in S1-CST neurons

We found that the majority of S1-CST neurons expressed mRNA encoding for the neuropeptide CCK. CCK expression in CST neurons suggests that they do not only release glutamate but likely also release CCK, which may modulate processing at the spinal level. CCK activates two distinct receptors (CCK-A and CCK-B) that are both expressed in the adult rodent spinal cord (Lein ES et al. 2007; Kim J et al. 2009; Haring M et al. 2018). In particular CCK-B has been reported to exert antagonizing effects on morphine-induced analgesia (Wiesenfeld-Hallin Z et al. 1999; Coudore-Civiale MA et al. 2000; Kovelowski CJ et al. 2000; Wiesenfeld-Hallin Z et al. 2002). However, CCK release has also been linked to anti-nociceptive effects (Roca-Lapirot O et al. 2019). At the level of the spinal cord, CCK is released from multiple sources, including local CCK neurons (Haring M et al. 2018), terminals of peripheral sensory neurons (Hokfelt T et al. 1994) and possibly from supraspinal sites (e.g. S1, anterior cingulate cortex, the thalamus, and the RVM (Fig. 1)). The release from different sites may contribute to the opposing effects or facilitate effects mediated by CCK.

### Molecular heterogeneity of S1-CST neurons

The results presented in this study suggest that S1-CST neurons are not a homogenous set of neurons but can be subdivided into subsets of neurons that differ in their molecular signature, as is the case for other projection neurons (Gerfen CR et al. 2013; Kim EJ et al. 2015; Xu NL 2020). The great majority, perhaps even all, of CST neurons in S1 express CCK, in addition to other well-established markers of cortical layer 5 neurons such as Ctip2 and ER81 (Arlotta P et al. 2005; Molyneaux BJ et al. 2007). Additionally, other genes such as the nuclear receptors *ROR*α (48.2%) or *Nurr1* (23.5%) are only expressed in subsets of CCK^+^ CST neurons. RORα and Nurr1 are nuclear receptors capable of driving specific transcriptional programs, suggesting that the subpopulations marked by the expression of the respective genes can be distinguished by a number of molecular markers. Until recently projection neurons within a cortical layer were considered as a rather homogenous population. However, in line with our findings recent RNAseq single cell data suggest a heterogeneous composition that is at least in part dependent on the target area (Xu NL 2020). Our data suggests that even cortical projection neurons that target the same CNS area (i.e. the lumbar dorsal horn of the spinal cord) are molecular heterogeneous. It is possible that this molecular heterogeneity translates into a functional heterogeneity of S1-CST neurons, comparable to what has been observed for other populations of cortical output neurons (Kim EJ et al. 2015; Klingler E et al. 2019).

### Presynaptic input to S1-CST neurons

In contrast to previous studies, we specifically targeted lumbar spinal cord projecting CST neurons in S1, thus enabling circuit analysis at an unprecedented resolution. Taking advantage of this, we were able to conduct retrograde monosynaptic tracing with rabies virus. Our experiments revealed that S1-CST neurons receive direct intracortical input from layer 2/3 pyramidal neurons as well as from inhibitory PV and to a lesser extend from NPY and SOM interneurons. Contrary to what has been suggested by previous studies (Tanaka YH et al. 2011; Cichon J et al. 2017) we found only little input from SOM interneurons. We see two possible explanations: First, the previous studies did not directly address direct connections between SOM neurons and specifically CST neurons in layer 5 of S1 and it is possible that different subsets of layer 5 pyramidal neurons receive distinct patterns of input (Anderson CT et al. 2010; Xu NL 2020). Second, we have previously shown that rabies-based retrograde transsynaptic tracing can be biased and does not always detect all inputs (Albisetti GW et al. 2017). We also identified direct input from sensory relay neurons in the VPL of the thalamus. This nucleus receives input from the postsynaptic dorsal column, the direct dorsal column pathway and the spinocervical tract that are known to propagate tactile information from the periphery to the brain (Abraira VE and DD Ginty 2013). Our data therefore indicates that CST neurons in S1 integrate direct ascending spino-thalamic sensory information with intra cortical input from layer 2/3 pyramidal neurons and local input from inhibitory interneurons. This is in agreement with the long-held view that sensory information relayed via the thalamus to the cortex is pre-processed by propagation from cortical layer 4 to layer 2/3 neurons, before arriving at the main output neurons of the somatosensory cortex located in the layer 5 (Gilbert CD and TN Wiesel 1979; Harris KD and GM Shepherd 2015). Moreover, it is also in agreement with *in vivo* evidence indicating the existence of direct connections between the thalamus and layer 5 neurons (Constantinople CM and RM Bruno 2013). The authors of the latter study suggested however that intracortical input might not be necessary for sensory evoked activity in layer 5 neurons. Bridging these opposing views, Quiquempoix et al. (Quiquempoix M et al. 2018) provided evidence that layer 2/3 pyramidal neurons primarily play a major role in tuning/amplifying sensory evoked responses in cortical output neurons of layer 5. Our data suggests that this model probably applies to CST neurons in layer 5 of S1.

### Non spinal target sites of CST neurons

When analysing the output of S1-CST neurons, we found evidence that, although they mainly project through the CST onto the spinal dorsal horn, they also send collateral branches to the dorsal striatum and thalamic nuclei and tectal areas. This extends, confirms and further specifies previous experiments in which CST collaterals were observed when all spinally projecting neurons were labelled (Wang X et al. 2017; Wang Z et al. 2018). Our data therefore indicate that, while their main target is the spinal dorsal horn (Ueno M et al. 2018), S1-CST neurons also contribute to cortico-striatal projections and may thereby influence goal-directed behaviours (Wang X et al. 2017). In addition, collaterals in the posterior thalamic nuclei are likely to influence thalamic integration of sensory inputs, possibly to adjust cortical responses to predicted versus unpredicted sensory signals (Groh A et al. 2014; Casas-Torremocha D et al. 2017).

### Spinal target neurons and functional implications on nociceptive signalling

Our anterograde tracing experiments revealed that CCK^+^ S1-CST neurons target different populations of spinal interneurons, with excitatory and inhibitory phenotypes being about equally prevalent among dorsal horn S1-CST target neurons. These results are also in line with findings from Ueno et. al, who showed that S1 and M1-derived CST neurons contact distinct populations of spinal interneurons (Ueno M et al. 2018). In addition, they provided evidence that the spinal neurons targeted by S1-CST neurons are involved in skilled movements. In another recent study, Liu et al. (Liu Y et al. 2018) observed changes in light touch sensitivity after ablation or silencing of S1-CST neurons in naïve and neuropathic states and proposed a neuropathic pain promoting role of S1-CST neurons. We found that a large portion of the inhibitory target neurons were glycinergic neurons and the majority of the excitatory neurons expressed c-Maf, a transcription factor necessary for the development of deep dorsal horn interneurons (Hu J et al. 2012). We have previously demonstrated that glycinergic dorsal horn neurons are an integral part of the spinal gate that controls spinal pain and itch relay (Foster E et al. 2015) and that activation of these neurons exerts a strong analgesic effect. Conversely, deep dorsal horn excitatory neurons have been linked not only to fine motor control (Bourane S et al. 2015; Ueno M et al. 2018) but also to chronic pain states (Peirs C et al. 2015; Cheng L et al. 2017; Liu Y et al. 2018). The c-Maf^+^ neurons identified in the present study are likely a subset of dorsal horn CCK^+^ excitatory interneurons (Haring M et al. 2018) and CST mediated activation of CCK^+^ spinal interneurons has strongly been linked to chronic pain states (Liu Y et al. 2018). Interestingly, we found no direct input of CST neurons onto PKCγ neurons. PKCγ neurons have also been strongly linked to neuropathic pain symptoms (Malmberg AB et al. 1997; Martin WJ et al. 2001) and they are also a subset of CCK^+^ spinal interneurons albeit different from the c-Maf^+^ interneurons. Our results therefore suggest that CST neurons could promote chronic pain states via engaging the c-Maf^+^ subset of CCK^+^ spinal interneurons. Conversely, it is likely that CST neurons can also have an analgesic function by stimulating deep dorsal horn glycinergic neurons.

### Outlook - Intersectional targeting of CST neurons

Most studies of the CST focused on its input to the ventral horn, its function in fine motor control (Wang X et al. 2017; Ueno M et al. 2018; Wang Z et al. 2018), and its role in the generation of deficits after spinal cord injury and their recovery (Steward O et al. 2008; Fry EJ et al. 2010; Jin D et al. 2015). Our data and the study by Liu et al. indicate that S1-CST target neurons also play a role in nociceptive signalling (Liu Y et al. 2018). Activation of all CST neurons in optogenetic or chemogenetic experiments would activate both inhibitory and excitatory spinal interneurons with potentially opposing effects on nociceptive signalling. Although unproven, it is well possible that different subtypes of spinal S1-CST target neurons (such as excitatory and inhibitory spinal interneurons) are innervated by distinct subsets of S1-CST neurons. We have shown that CST neurons in S1 are not a homogenous population. The intersectional targeting strategies presented here allow further dissecting of the function of potential CST subpopulations. For example, applying the same intersectional targeting strategies to RORα^Cre^ mice will uncover which neuronal subtypes are targeted by the RORα^+^ subset.

Alternatively, retrograde tracing initiated at the level of a specific spinal target populations identified by us and others (Abraira VE et al. 2017; Ueno M et al. 2018) may be used in order to identify target-specific sub-networks and functionally interrogate subpopulations of S1-CST neurons. Self-inactivating rabies virus such as those recently developed by Ciabatti et al. (Ciabatti E et al. 2017) may be suitable to functionally manipulate the S1-CST neuron population that innervates the respective spinal target populations. Finally, our study highlights the importance of spatially restricted and intersectional manipulation of either CST or spinal neurons that express a given marker gene, as several of the genes that we and others found expressed in CST neurons are also present in spinal cord neurons (e.g. *CCK, RORα* or *Pkcγ*).

## Supplementary figures

**Fig. S1.**
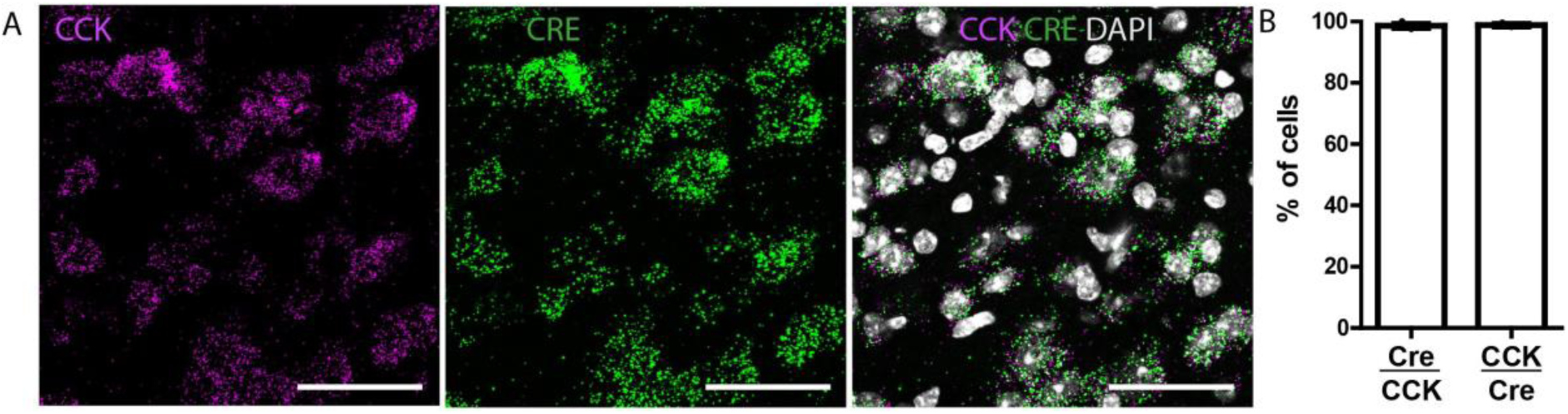
Overlap between CCK and Cre mRNA expression in CCK^Cre^ mice: (A) Double ISH showing the colocalization of Cre with CCK mRNA. (B) Quantification of (A): 98.6 ± 1.2% CCK positive neurons are Cre positive and 98.8 ± 0.7% Cre positive neurons are CCK positive (n = 3 mice; 1087 neurons). Error bars represent ± SEM. Scale bars: 50 µm.

**Fig. S2 (related to Fig. 3):**
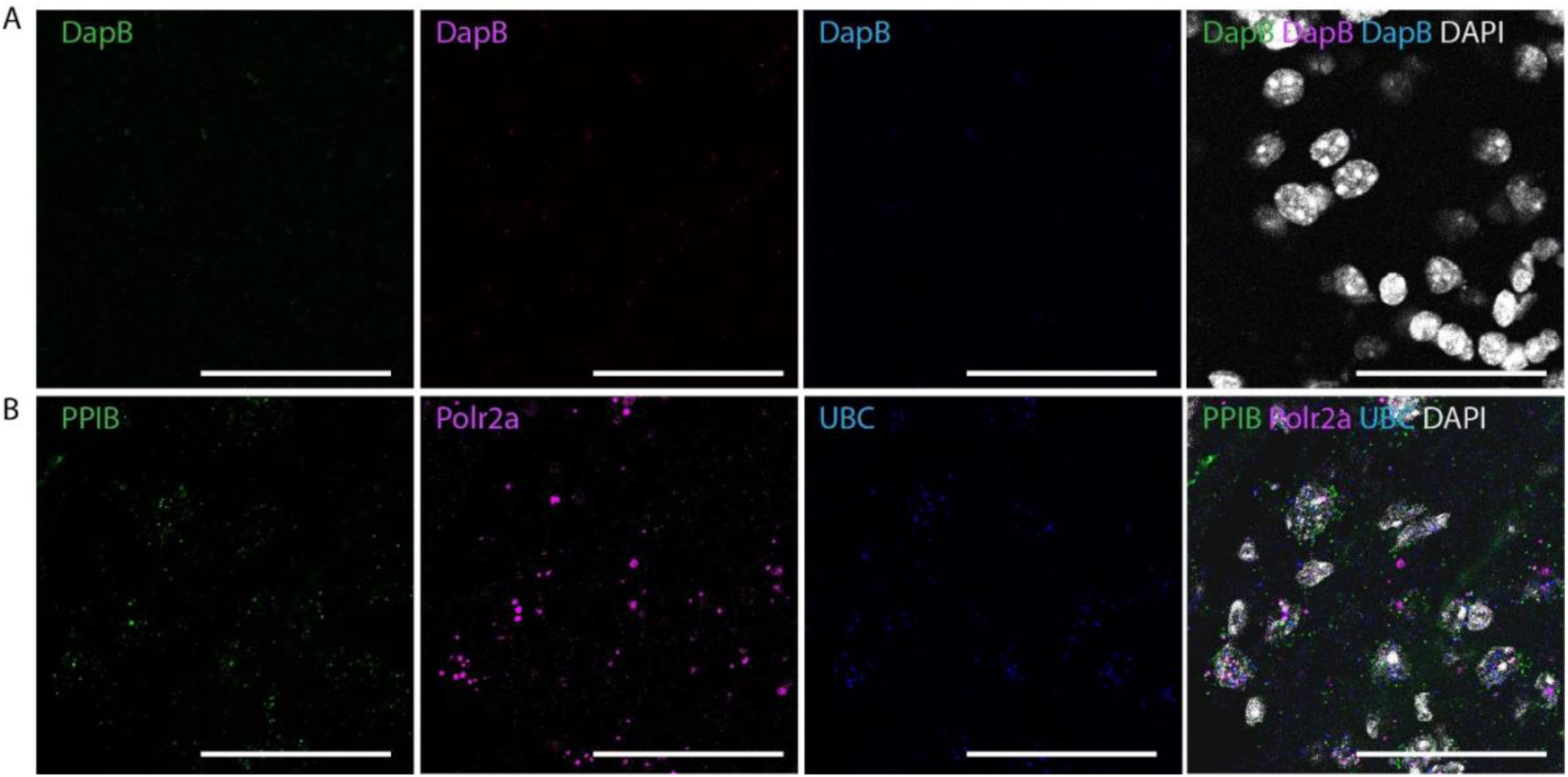
Controls for the multiplex ISH in GFP-labelled CCK^Cre^ neurons. (A) Triple ISH negative control showing signals for the marker *DapB*. (B) Triple ISH positive control showing signals for the markers *PPIB, Polr2a* and *UBC*. (n = 3 mice). Scale bars: 50 µm.

**Fig. S3 (related to Fig. 4):**
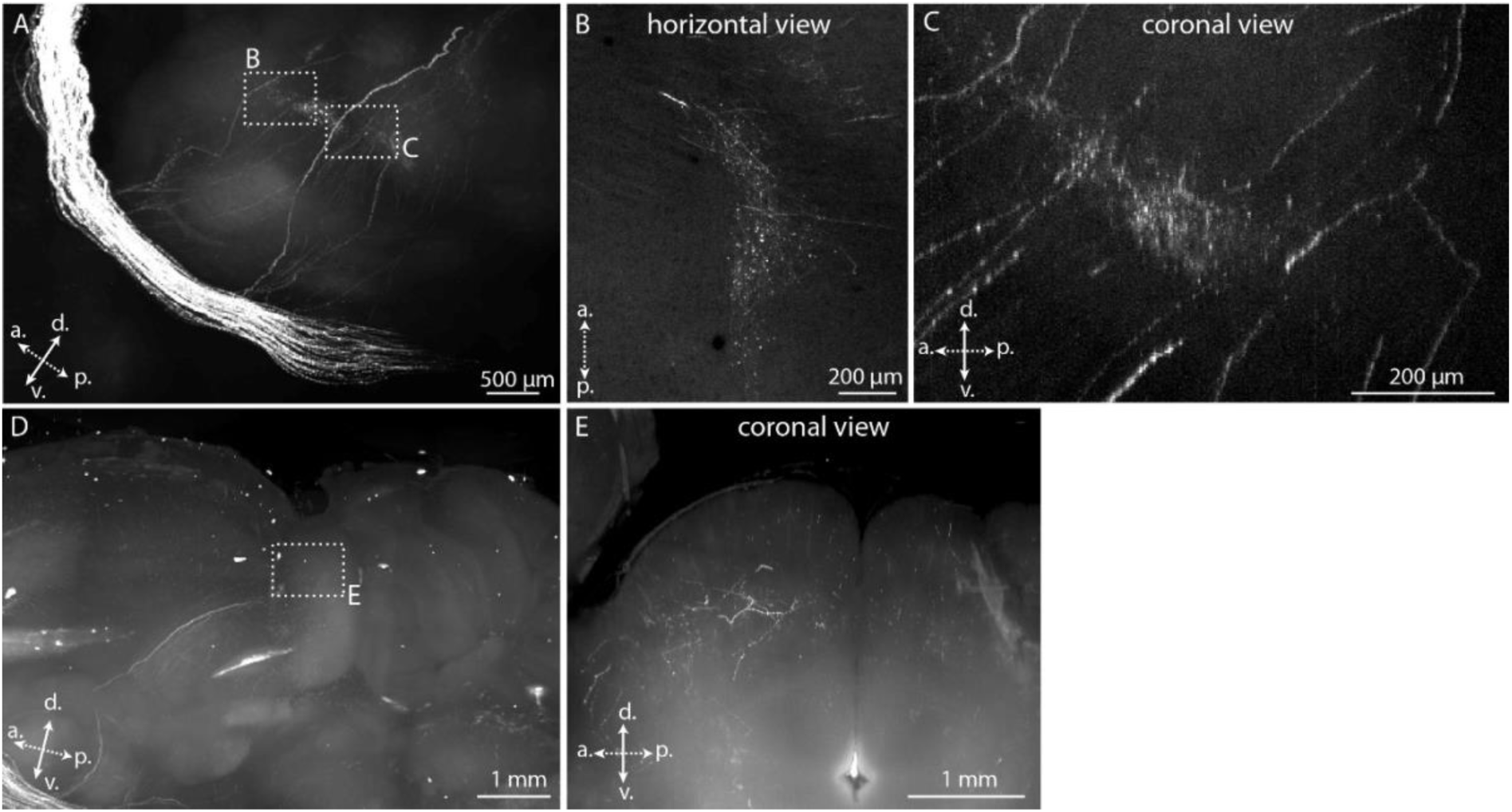
Detail of the CST collaterals in the thalamus. (A) Collaterals branching of from the main CST towards the thalamus (Fig. 4C), showing 2 termination areas (B) and (C). (C) Collaterals terminating in tectal areas. (E) Coronal view of inset from (D). (n = 3 mice). a.: anterior, p.: posterior, d.: dorsal, v.: ventral.

**Fig. S4 (related to Fig. 6):**
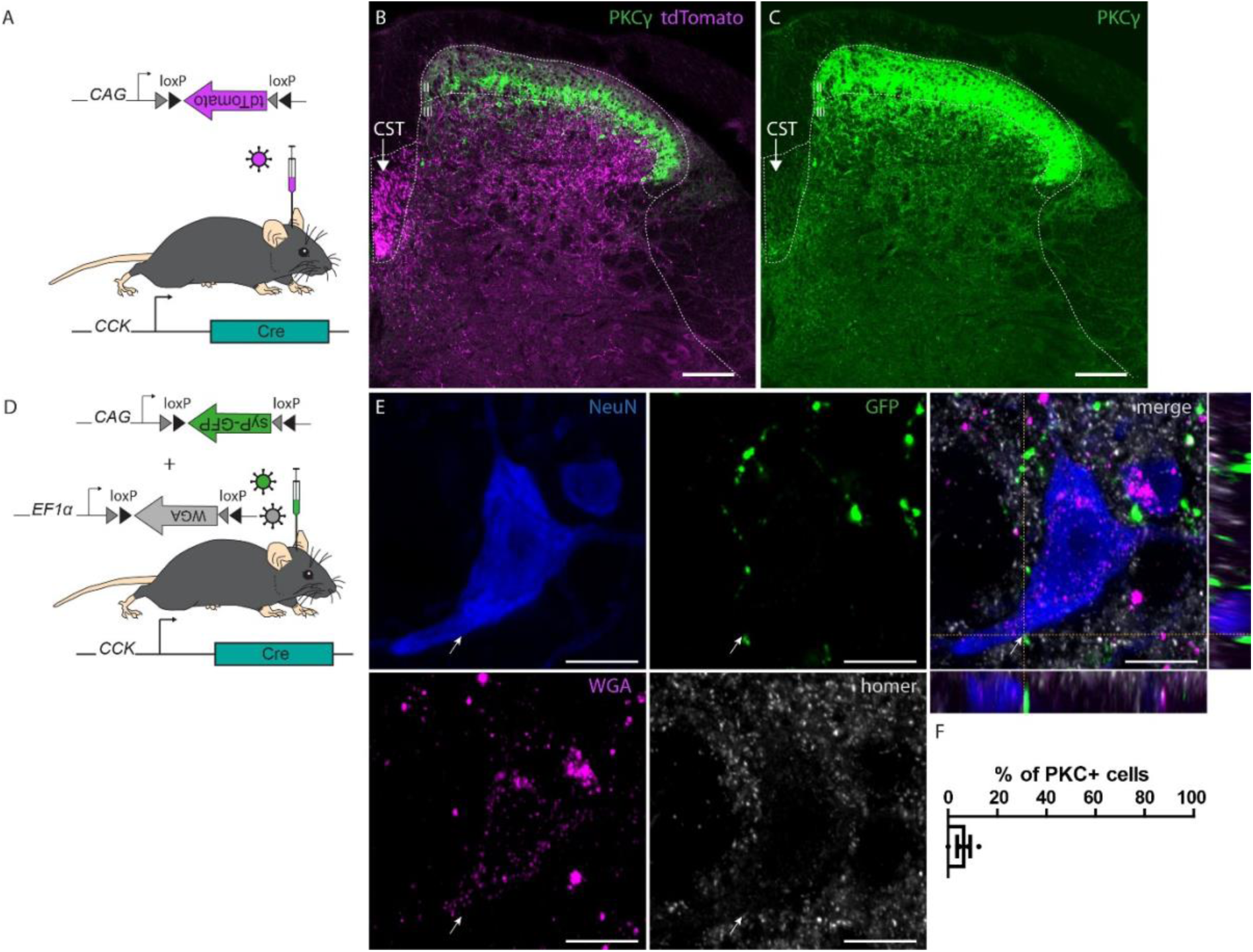
Localisation of S1-CST neurons terminals in the spinal cord. (A) rAAVs carrying a Cre-dependent tdTomato were injected in the S1hl of CCK^Cre^ mice. (B) Labelling of the CST in the dorsal funiculus of the spinal cord (same sample as in Fig. 6A), contralateral to the brain injection site. CST terminals are preferentially located below the laminae II-III border marked by PKCγ immunoreactivity. (C) The CST is also labelled by PKCγ immunoreactivity. (D) rAAVs carrying a Cre-dependent WGA and a Cre-dependant synaptophysin-eGFP fusion protein transgenes were injected in S1hl of CCK^Cre^ mice. (E) Representative example of a WGA^+^ neuron in close proximity to a GFP^+^ presynaptic terminals of S1-CST neurons after co-injection of rAAVs carrying a Cre-dependent WGA and a Cre-dependant synaptophysin-eGFP fusion protein transgenes in S1hl of CCK^Cre^ mice (from a different animal as the example presented in Fig. 6, n =4). (F) Quantification of the number of PKCγ^+^ neurons receiving direct contacts from eGFP^+^ synaptic terminals (n = 4 mice; 98 neurons). CST: corticospinal tract Error bars represent ± SEM. Scale bars: B-C: 100 µm; E: 10 µm.

## Acknowledgments

The work has been supported through the Clinical Research Priority Program of the University of Zurich (CRPP Pain). N. Frezel has been supported through a Contrat Doctoral Specifique pour Normaliens (CDSN) grant awarded for a joint PhD at the University of Zürich and the Institute of Biology of the École Normale Supérieure (IBENS), Paris Sciences et Lettres Research University (PSL), Paris 75005, France. E. Platonova has been supported through the Technology Platform commission of the University of Zürich. F. Helmchen was supported by an European Research Council (ERC) Advanced Grant (project no. 670757). R. Kastli and T. Karayannis were supported by grants from the European Research Council (ERC, 679175, T.K), the Swiss National Science Foundation (SNSF, 31003A_170037, T.K) and the Swiss Foundation for Excellence in Biomedical Research (R.K and T.K). We want to thank Jean-Charles Paterna (Viral Vector Facility, UZH, Zurich, Switzerland) for the production of viral vectors.

## Contributions

N.F carried out the experiments, E.P, J.M.M, U.Z., R.K, T.K. F.H. and F.F.V contributed to the CLARITY experiments and helped with the analysis. N.F., H.W and H.U.Z wrote the manuscript. H.W and H.U.Z. supervised the project. All authors have read and commented on the manuscript.

## Abbreviations

CST: corticospinal tract
rAAV: recombinant adeno-associated virus
RVM: rostral ventromedial medulla
S1: primary somatosensory cortex
S1hl: hindlimb area of S1
S1-CST: corticospinal tract originating in S1
7N: facial nuclei
ACC: anterior cingulate cortex
MnR: median raphe nucleus
PAG: Periaqueductal grey
S1hl: somatosensory cortex
hindlimb area.VPL: ventral posterolateral nucleus of the thalamus
WGA: wheat germ agglutinin
Cpu: dorsal striatum.

